# The HNF4A Q164X Mutation Impairs Transcriptional Activation in Vitro but Its Heterozygosity Suppresses Liver Tumorigenesis in Vivo

**DOI:** 10.1101/2025.10.28.685175

**Authors:** Dawid Winiarczyk, Hossein Khodadadi, Effi Haque, Piotr Poznański, Mariusz Sacharczuk, Hiroaki Taniguchi

## Abstract

Hepatocyte nuclear factor 4 alpha (HNF4A) is a master regulator of hepatic differentiation and metabolism. Here, we identify and characterize a truncating Q164X mutation that impairs HNF4A transcriptional activity in vitro and causes embryonic lethality when homozygous. Functional assays revealed that the Q164X protein retains nuclear localization but exhibits severely reduced DNA binding and transcriptional activation. CRISPR-generated Q164X mice showed no viable homozygotes, confirming the essential role of HNF4A in early embryogenesis. Unexpectedly, heterozygous Q164X mutants displayed reduced liver tumorigenesis following diethylnitrosamine and high-fat diet treatment, despite downregulation of HNF4A target genes such as *ApoB* and *Hnf1a*. These results suggest that partial HNF4A deficiency may trigger compensatory metabolic networks that protect against carcinogenic stress. Collectively, our study establishes Q164X as a loss-of-function HNF4A mutation with paradoxical tumor-suppressive effects in vivo.

## Introduction

Hepatocyte nuclear factor 4 alpha (HNF4A) is a liver-enriched nuclear receptor that functions as a master regulator of hepatocyte differentiation and metabolic homeostasis. Through its DNA-binding (zinc-finger) and ligand-binding domains, HNF4A orchestrates transcriptional programs governing glucose, lipid, and amino acid metabolism while repressing proliferative and epithelial–mesenchymal transition (EMT) genes^1–9^. Loss or reduction of HNF4A expression has been repeatedly observed in hepatocellular carcinoma (HCC) and is associated with poor differentiation, aggressive tumor behavior, and unfavorable clinical outcomes^10^. Collectively, these findings position HNF4A as a tumor suppressor and a central determinant of hepatic identity. Among known pathogenic variants, nonsense and truncating mutations affecting the C-terminal hinge or ligand-binding regions can severely impair transcriptional activation by disrupting dimerization and cofactor interactions^11–13^. The Q164X mutation, originally identified in Japanese liver cancer patients^14,15^, though its functional significance has not been experimentally verified, introduces a premature stop codon within this region, yielding a truncated protein that retains the zinc-finger domain but lacks critical structural elements required for interactions with transcriptional coactivators such as CBP/p300^12^ and PGC-1α^16^. Consequently, this mutation is expected to uncouple DNA binding from downstream transcriptional activation, thereby functionally inactivating HNF4A’s tumor-suppressive axis.

Recent evidence, including our present findings, indicates that the integrity of HNF4A’s zinc-finger domain plays an essential role in preventing liver cancer progression^14,17^. Our study further reveals that while the Q164X mutation abolishes the transcriptional function of HNF4A *in vitro*, its heterozygous presence in mice unexpectedly leads to reduced tumor development compared with WT animals, despite the downregulation of HNF4A target genes such as *ApoB* and *HNF1A*. These observations suggest that HNF4A inactivation in liver cancer is not solely attributable to reduced expression levels but can also result from structural mutations that decouple its transcriptional control. Interestingly, while homozygous HNF4A mutations are embryonically lethal, heterozygosity for Q164X appears to suppress, rather than promote, tumor progression. Understanding this paradoxical behavior will be an important focus of future studies and supports the notion that single-allele mutations in HNF4A may not necessarily drive hepatocarcinogenesis.

## Materials and Methods

### Animals

All animal experiments were conducted at the Institute of Genetics and Animal Biotechnology, Polish Academy of Sciences (IGAB PAS). Animals were housed in a temperature-controlled facility under a 12-hour light/12-hour dark cycle (lights on from 06:00 to 18:00). All procedures were performed in accordance with the Polish Governmental Act for Animal Care and were approved by the II Local Ethical Committee for Experiments on Animals in Warsaw (Approval No. WAW2/006/2023).

### Generation of Q164X Mouse Model

The Q164X nonsense mutation (transcript *Hnf4a*-201, ENSMUST00000018094.13) was introduced into the *Hnf4a* locus of C57BL/6JRj mice using CRISPR/Cas9-mediated homology-directed repair. A modified TrueGuide™ sgRNA (Invitrogen) targeting exon 4 and a single-stranded DNA repair template (EXTREmers, Eurofins Genomics; purified with KAPA beads) containing the desired mutation(5′CAGCACGCGGAGGTCAAGCTACGAGGACAGCAGCCTGCCCTCCATCAACGCGCTCCTGCAGGCAGAGGTTCTGTCTCAGTAGgtaccaaggaatcctctttactacccaggctcccttccaggggaaatcacttcttctacttctgtctc-3′) were co-injected with Cas9 mRNA into fertilized C57BL/6JRj zygotes.

Resulting embryos were transferred into pseudopregnant females, and founder mice were screened by PCR using primers Fw: 5′-catagCTGTCCAAAATGAGC-3′ and Rv1: 5′-ctatccatgtggacctccag-3′ (shorter amplicon) or Rv2: 5′-gccacctgctaccatgttac-3′ (longer amplicon). PCR products were confirmed by Sanger sequencing using the same primer set (primarily Rv1). The mouse model was generated with the Genome Engineering Unit of the International Institute of Molecular and Cell Biology in Warsaw.

### Diethyl-nitrosamine and High-fat diet treatment

We conducted two independent experiments. In the first, liver samples were collected from WT and HNF4A Q164X/WT mice at 6 months of age under non-stress conditions. In the second, to promote tumorigenesis, mice were treated with diethylnitrosamine (DEN) and a high-fat (HF) diet. At two weeks of age, male mice received a single intraperitoneal (IP) injection of Diethyl-nitrosamine (DEN) (25 mg/kg body weight). Both HNF4A Q164X/WT and wild-type (WT) mice were treated under the same conditions. Six weeks after DEN injection, both groups were fed a high-fat diet (HFD) containing 20% fat and 1% cholesterol (Altromin, Germany). The experimental setup consisted of two groups: WT (n = 7) and HNF4A Q164X/WT (n = 7) males, both subjected to the HFD for 40 weeks to induce obesity and promote liver tumorigenesis. At the end of the experiment, mice were sacrificed, and tissue samples were collected for subsequent analyses.

### Tissue preparation

After confirmation of death manifested by lack of corneal reflex, breath, heart beating and absence of response to hind paw pinching, two fragments of left lobe of the liver were collected and fixed in 4% PFA for additional 24h in 4℃. After that, one fragment was dehydrated through ascending ethanol gradient (30%, 50%, 70%, 80%, 95%, 99.8%), cleared in three changes of xylene (15 minutes each change) and embedded in paraffin blocks. Second fragment was cryoprotected in ascending sucrose gradient (10%, 20%, 30%), sample was kept in each solution one hour and then overnight in 30% sucrose solution. Positive saturation was confirmed after tissue fell down to the bottom of the probe. After that, tissue was frozen at -80℃ in the optimal cutting temperature compound (OCT). Paraffin blocks were sectioned with microtome (Hyrax M25, Zeiss, Germany) for 3µm thick sections. Cryosectioning was done with cryostat (Jung CM1800, Leica, Germany) for 10µm sections at -17℃. Prior to cryosectioning embedded tissue was transferred to cryostat chamber for 1h to equilibrate temperatures. All sections were placed on gelatin-coated microscopic slides to prevent sections detaching during staining. Sodium pentobarbital was bought from Biowet Puławy (Puławy, Poland). Phosphate-buffered saline (PBS) as well as Shandon™ Cryomatrix was provided by ThermoFisher Scientific (Warsaw, Poland). Harris hematoxylin, eosin solution, oil red O stock solution, sucrose, paraffin wax, paraformaldehyde powder and DPX mounting medium were purchased from Sigma-Aldrich (Saint Louis, MO, USA). Ethanol, isopropanol, xylene was supplied by Warchem Sp. z.o.o. (Zakręt n/Warsaw, Poland).

### Hematoxylin and eosin staining

Hematoxylin and eosin staining (H&E) was done on 3µm thick paraffin sections according to standard protocol at room temperature. Briefly, sections were deparaffinized in two changes of xylene, 15 minutes each and rehydrated in descending ethanol gradient (two changes of 99.8%, 95%, 80%, 70%, 50%, 30%), 10 minutes each. Then, slides were washed briefly in two changes of distilled water. After that, slides were stained in hematoxylin solution for 5 minutes, washed in distilled water and immersed in eosin solution for 15 seconds, followed by distilled water washing. Finally, sections were dehydrated in ascending ethanol gradient (30%, 50%, 70%, 80%, 95% and two changes 99.8%), 10 minutes each, cleared in xylene (two changes for 15 minutes each) and mounted with DPX.

### Oil red O staining

Oil red O staining (ORO) was done on 10 µm thick cryosections according to well-established protocol. Briefly, slides were washed in distilled water for 5 minutes, followed by 60% isopropanol rinse. Immediately, slides were placed in freshly prepared ORO working solution for 15 minutes, followed by wash in 60% isopropanol solution. Then, lightly counterstained with hematoxylin, obtained by 30 seconds incubation. To remove excess hematoxylin, prior to mounting, slides were washed in distilled water. Coverslips were mounted with glycerine jelly mounting medium and stored in 4°C for further examination.

### Cell Culture

HEK293, CV-1, and HuH7 liver cancer cells were cultured in Dulbecco’s Modified Eagle Medium (DMEM; high glucose, Biowest, France) supplemented with 10% fetal bovine serum (FBS; EURx, Poland), 4.5 g/L glucose, 100 µg/mL streptomycin, and 100 U/mL penicillin. The cells were maintained in a humidified incubator at 37 °C with 5% CO_2_.

### RNA Extraction and cDNA Synthesis

Total RNA was extracted from HuH7 cells and mouse liver tissue. The isolated RNA was used as a template for cDNA synthesis using the NG dART RT kit (EURx, Gdańsk, Poland), following the manufacturer’s instructions. The reactions were carried out in a GeneAmp® PCR System 9700 thermal cycler (Applied Biosystems, Waltham, MA, USA) under the following conditions: 50 °C for 60 min, 85 °C for 5 min, and cooling to 4 °C.

### Quantitative Real-Time PCR

Gene expression levels identified from transcriptome sequencing were validated by Quantitative real-time PCR (qPCR) for selected genes using the RT HS-PCR Mix SYBR® C (A&A Biotechnology, Gdynia, Poland), according to the manufacturer’s instructions. Primer sequences (purchased from Sigma-Aldrich, St. Louis, MO, USA) are Gapdh (FW: TGACCTCAACTACATGGTCTACA, RV: CTTCCCATTCTCGGCCTTG), Hnf1a (FW: GTCGAACATCCAGCACCTG, RV: GGCCATCTGGGTGGAGATAA), Apo B (FW: GGCAAGCATGAACAGGACAT, RV: AAGTTTGTGCACCACTCAGC). qPCR reactions were performed using the LightCycler® 480 Instrument (Roche, Mannheim, Germany).

### Western blotting

A total of 5 × 10^5^ HEK293 cells were seeded in 6-well plates and transfected with 2 µg of each HNF4A expression plasmid for 48 h using Lipofectamine 3000 (Thermo Fisher Scientific), according to the manufacturer’s instructions. Nuclear protein concentrations from HNF4A WT and mutant overexpression cells were determined using the Pierce BCA Protein Assay Kit (Thermo Fisher Scientific, Waltham, MA, USA). Protein molecular weights were estimated using Precision Plus Protein Western C Standards (Bio-Rad, Hercules, CA, USA). A total of 10 µg of each protein sample was loaded onto an SDS– polyacrylamide gel, separated, and transferred to a PVDF membrane (Merck Millipore, Burlington, MA, USA) by wet transfer. The membranes were blocked with 5% skim milk for 15 min and then incubated with primary and secondary antibodies. The blot was incubated overnight at 4 °C with a mouse monoclonal anti-FLAG antibody (1:5000, Sigma-Aldrich, St. Louis, MO, USA) diluted in 1% skim milk and 0.1% PBST, followed by incubation for 1 h at room temperature with an HRP-conjugated goat anti-mouse IgG (1:5000, Sigma-Aldrich) in the same buffer. The proteins were visualized using an ECL Western Blotting Detection System (Amersham, Illinois, USA) and imaged with the ChemiDoc XRS+ System (Bio-Rad).

### Cell Proliferation Test (WST-1 Assay)

The effect of the selected HNF4A mutation on cell proliferation and growth was evaluated using the Cell Proliferation Reagent WST-1 (Roche, Mannheim, Germany). HuH7 cells were electroporated with WT or mutant HNF4A constructs and seeded into 24-well plates. After 24 h and 48 h of culture, cell proliferation was assessed according to the manufacturer’s instructions. Absorbance was measured using a Synergy LX Multi-Mode Reader (BioTek, Winooski, VT, USA).

### Immunofluorescence

Cells were fixed with 4% paraformaldehyde for 15 min at room temperature. After washing with PBS containing 0.1% Tween-20 (PBST), the cells were blocked with 1% skim milk for 20 min at room temperature. The cells were then rinsed once with 0.1% PBST, permeabilized with 0.5% PBST for 10 min, and incubated with a mouse monoclonal anti-FLAG antibody, followed by extensive washing and incubation with an Alexa Fluor 546– conjugated anti-mouse IgG secondary antibody (Thermo Fisher Scientific, Waltham, MA, USA) for 1 h. After three additional washes with PBST, the nuclei were stained with DAPI, and fluorescence images were captured using a fluorescence microscope. Nikon A1R confocal microscope (Tokyo, Japan). The images were captured using the NIS-Elements package. Confocal images were analyzed using IMARIS 6.0.1 software (Bitplane AG, UK).

### Electrophoretic Mobility Shift Assay (EMSA)

Double-stranded probes were prepared by annealing equimolar amounts of each sense and antisense oligonucleotide at 95 °C for 10 min, followed by gradual cooling to room temperature. The resulting double-stranded oligonucleotides were labeled with DIG-11-ddUTP using recombinant terminal transferase (20 U/mL) in a final reaction volume of 25 μL, according to the manufacturer’s instructions for the DIG Gel Shift Kit, Second Generation (Roche Applied Science, Mannheim, Germany).

EMSA was performed following the manufacturer’s protocol. Briefly, DNA-binding reactions were assembled using 5 μg of nuclear extract from cells expressing either wild-type or mutant HNF4A proteins. The extracts were incubated with the DIG-labeled probes in a DNA-binding buffer containing 1 μg poly(dI–dC) and 0.1 μg Poly-L-lysine, in a final volume of 20 μL. The HNF4A supershift assay, used to confirm the specific binding of HNF4A to the probe, has been previously described^14^.

### BGI RNA-Seq (Transcriptome) Sequencing

Total RNA was extracted from HuH7 cells 48h after transfection with the N-terminal p3XFLAG-CMV plasmid (Sigma-Aldrich, St. Louis, MO, USA) or plasmids containing WT and mutant HNF4A using the NucleoSpin RNA Kit (Macherey-Nagel, Düren, Germany). Changes in gene expression were analyzed using the DNBSEQ platform (BGI Genomics, Shenzhen, China). Bioinformatic analyses were performed using BGI’s proprietary software by Dr. Tom (BGI Genomics). Transcriptome analysis was also conducted on liver tissues collected from 6-month-old wild-type (WT) and HNF4A Q164X/WT heterozygous mice under non-stress conditions. Raw sequencing data are being deposited in GEO and will be made publicly available before journal submission.

### Cell Transfection and Luciferase Assays

For the luciferase assays, 2 × 10^4^ cells were seeded into 24-well plates. After 24 h, the cells were transfected with HNF4A WT and mutant plasmids together with an HNF4A reporter using Lipofectamine 3000 (Thermo Fisher Scientific, Waltham, MA, USA). Following 48 h of incubation, the cells were harvested, and luciferase activity was measured using the Luciferase Assay System (Promega, Madison, WI, USA) according to the manufacturer’s instructions, with luminescence detected on a Luminometer. All plasmid information used in this study was provided in reference^14^.

### Statistical Analysis

Data are presented as the mean ± standard error of the mean (SEM) from at least three independent experiments. Statistical significance was evaluated using either Student’s t-test or one-way ANOVA followed by Tukey’s post hoc test, where applicable. A *p*-value < 0.05 was considered statistically significant. GraphPad Prism was used for statistical analysis.

## Results

From an evolutionary perspective, the HNF4A Q164 residue analyzed in this study is highly conserved among various species, including humans, mice, cattle, zebrafish, and flies (Fig. 1A). The red asterisk indicates the position of Q164 within the alignment. The conserved domains among the analyzed species are highlighted in red, and the regions subjected to functional analysis in this study are 100% conserved across all examined species. Mutations occurring in such evolutionarily conserved elements are expected to exert a strong impact on protein function, although further studies are needed to clarify the precise mechanisms. Fig. 1B shows the location of Q164 within the human HNF4A protein, which is composed of a zinc-finger DNA-binding domain and a ligand-binding domain.

**Fig. 1.**
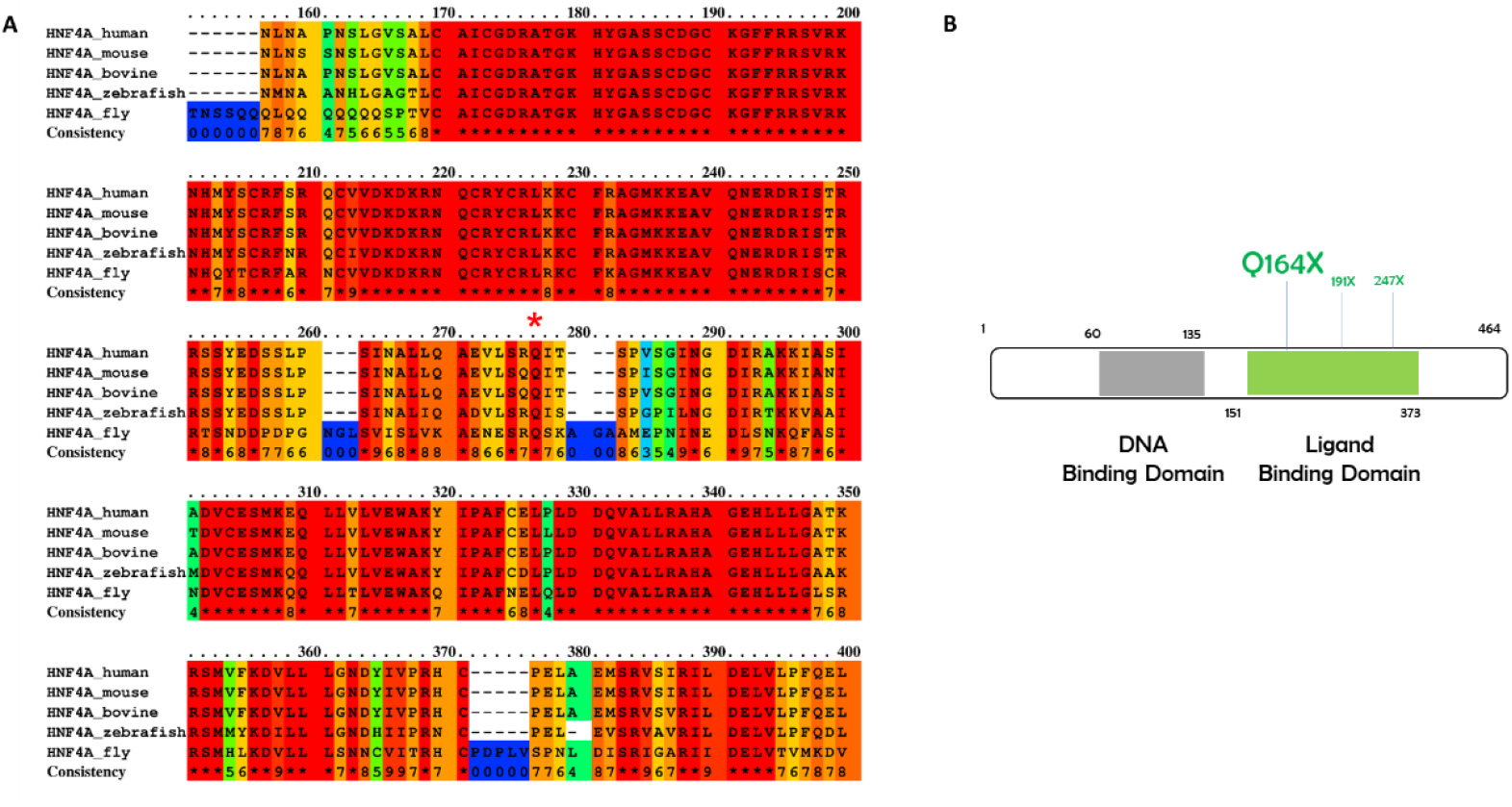
Evolutionary conservation and location of the HNF4A Q164X mutation. **(A)** Alignment of *HNF4A* amino acid sequences from human, mouse, bovine, zebrafish, and fly by PRALINE software. Highly conserved residues (100% conservation across species) are highlighted in red boxes. The location of the Q164X mutation in *HNF4A* is indicated by an asterisk. **(B)** Locations of the novel mutations are shown on the human HNF4A protein structure (DNA-binding domain, gray; ligand-binding domain, light green).

Immunofluorescent staining showed that both WT and mutant form of HNF4A Q164X were localized in the nuclei of HEK293 cells (Fig. 2A), indicating that these amino acid substitutions do not affect nuclear transport of the protein. However, the transcriptional activity of the HNF4A Q164X mutant was markedly reduced compared with WT (Fig. 2C), suggesting that this mutation interferes with DNA binding or coactivator interaction. To assess DNA-binding ability, electrophoretic mobility shift assays (EMSA) were performed using oligonucleotides derived from the HNF4A-binding elements of the *HNF1A* promoter. Consistent with its reduced transcriptional activity, the Q164X mutant displayed markedly diminished binding to both promoter elements compared with WT HNF4A, whereas protein expression levels were comparable among constructs (Fig. 2B). These findings indicate that the Q164X mutation compromises DNA binding rather than protein stability or localization. Functionally, both G79C and Q164X mutations attenuated the ability of HNF4A to suppress cell proliferation, consistent with loss-of-function characteristics of a tumor suppressor (Fig. 2D). Furthermore, RNA-seq analysis using HuH7 cells expressing WT or mutant HNF4A revealed that HNF4A WT upregulated 230 genes, whereas only 34 and 23 genes were upregulated in the G79C and Q164X mutants, respectively (Fig. 2E). Collectively, these results demonstrate that G79C and Q164X mutations cause a profound loss of HNF4A transcriptional function, primarily through impaired DNA binding rather than altered subcellular localization.

**Fig. 2.**
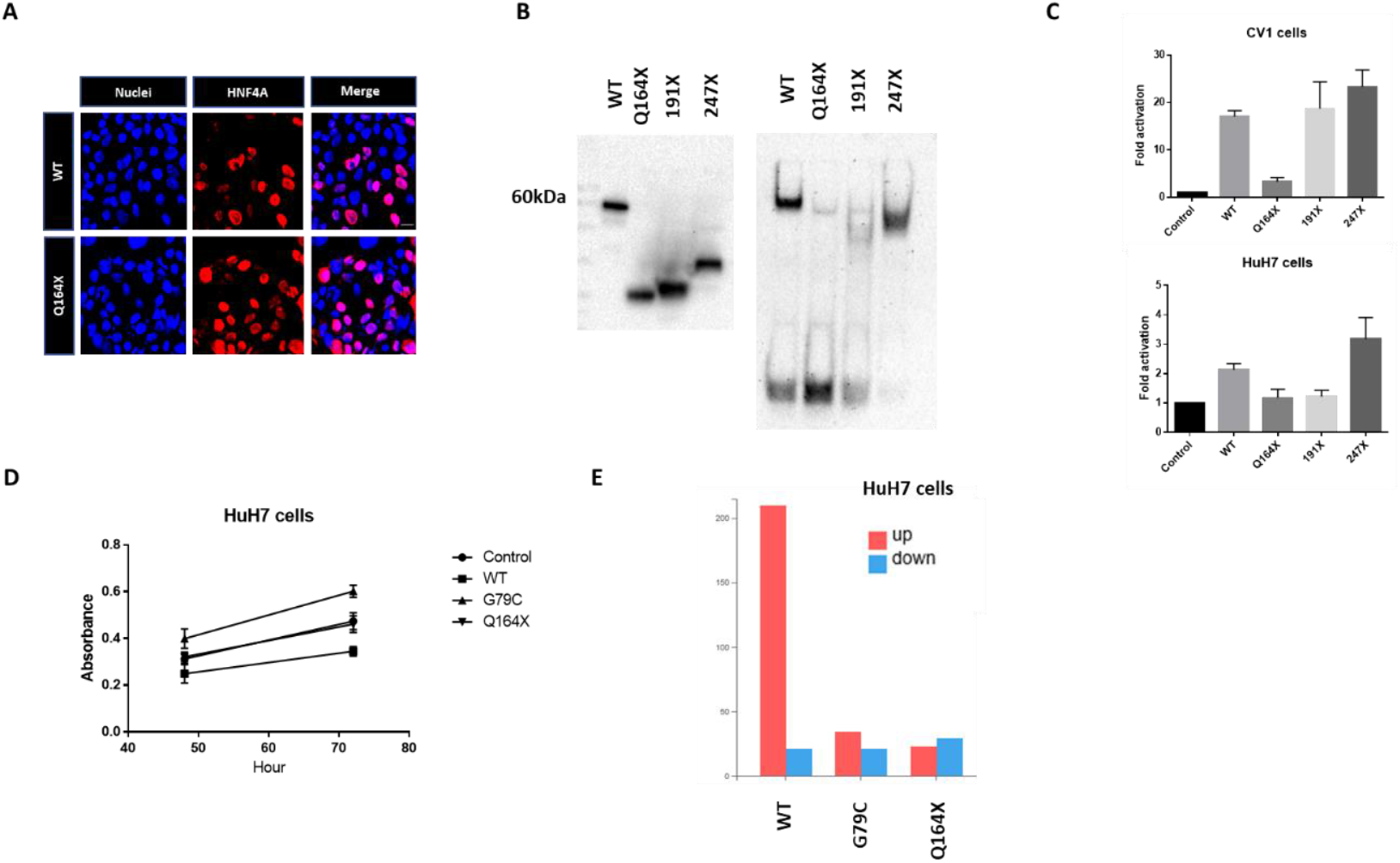
Functional characterization of HNF4A Q164X mutant in vitro. **(A)** Nuclear localization of HNF4A mutants using immunofluorescent staining Cellular localization of WT and mutant HNF4A was visualized in HEK293 cells. The nuclei were stained with DAPI. (**B)** Electrophoretic mobility shift assays were used to assess the binding of WT or mutated HNF4A nuclear proteins to a double-stranded oligonucleotide corresponding to the consensus HNF4A binding elements of the HNF1A promoter region. HEK293 cells were transfected with expression vectors encoding WT HNF4A or the indicated mutants. Western blot analysis showed that all proteins were expressed at comparable levels. **(C)** The ability of WT and mutant HNF4A to transactivate target promoters when overexpressed in CV-1 and HuH7 cells. The cells were co-transfected with the indicated luciferase reporters along with either an empty expression vector (serving as a control) or expression vectors (50 ng) for the indicated HNF4A proteins in 24-well culture plates. The bars indicate the fold activation of HNF4A WT and mutants (vs. control) on HNF4A target promoters. Promoter activity is presented as fold activation over control (mean ± SEM, *n* = 3). Three independent experiments were performed, each in duplicate, and the data represent the average of these experiments. (**D)** Effect of HNF4A mutant overexpression on cell proliferation in HuH7 cells (48-72h). A significant loss of HNF4A-dependent suppression of cell proliferation was observed in cells expressing the HNF4A Q164X and G79C mutants compared with control cells. **(E)** Effect of HNF4A overexpression on transcription in HuH7 cells. RNA-seq analysis revealed both downregulated and upregulated genes in HuH7 cells expressing WT, G79C, or Q164X HNF4A.

To investigate the *in vivo* effect of the Q164X mutation, we established a time- and cost-efficient method to evaluate the outcome of zygote electroporation combined with a cloning-free CRISPR/Cas9 system^18^. Using this approach, we introduced HNF4A mutations via ssDNA donors to generate the desired nucleotide changes. However, we found that this method efficiently introduced the targeted mutation into both alleles, resulting in embryonic lethality.

To overcome this, we replaced the Cas9 protein with Cas9 mRNA, allowing delayed Cas9 translation and thus postponing homologous-directed repair to later embryonic stages (e.g., the 2-cell stage). This modification enabled us to obtain heterozygous mutations. Using this optimized approach, we successfully generated HNF4A Q164X mutant mice (Fig. 3A). Interestingly, no homozygous pups were recovered (36 genotyped: 17 WT, 19 Q164X heterozygotes, and 0 Q164X homozygotes), a statistically significant deviation from the expected Mendelian ratio (*p* < 0.0003, assuming heterozygotes are viable). Consistent with previous reports showing that complete knockout of *HNF4A* leads to embryonic lethality, this result supports that the Q164X mutation causes a loss of function *in vivo*, in agreement with our *in vitro* data.

**Fig. 3.**
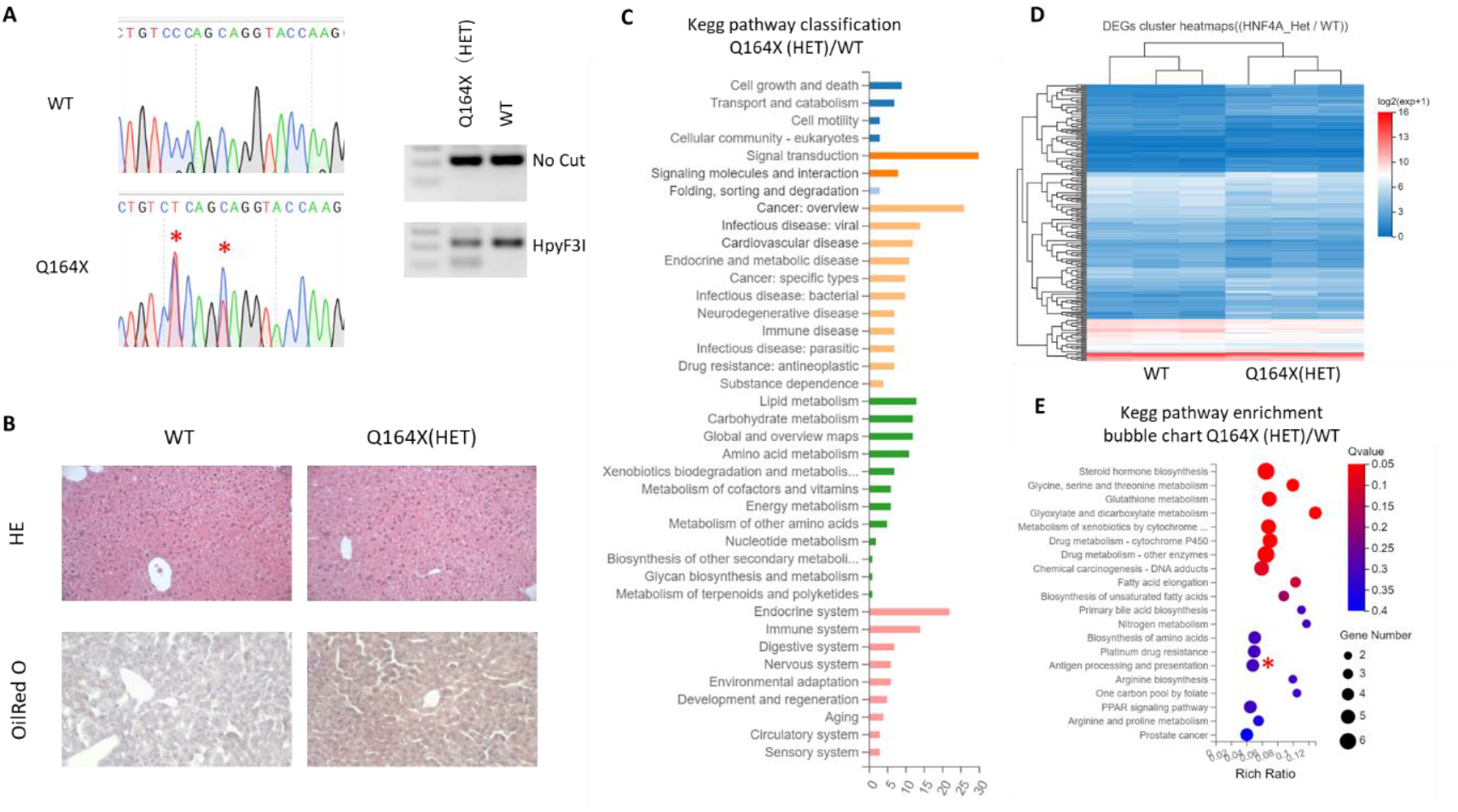
Generation and hepatic characterization of HNF4A Q164X mutant mice. **(A)** Direct sequencing and restriction enzyme digestion confirm the CRISPR-mediated generation of Q164X mutant mice. The newly introduced mutation is recognized by the HpyF3I enzyme and produces cleaved bands when the PCR product is digested. **(B)** Oil Red O staining reveals lipid accumulation in the livers of Q164X mice at 6 months of age. **(C-E)** RNA-seq analysis identified 230 genes significantly changed in heterozygous (Het) Q164X mice compared with WT controls. KEGG pathway analysis indicates disruption of lipid metabolic pathways in the livers of Q164X mice.

We next analyzed heterozygous Q164X mutant mice at six months of age. Body weight and liver-to-body weight ratios did not differ significantly between WT and heterozygous animals (not shown). Histological examination, however, revealed increased lipid accumulation in the livers of Q164X mutants, as shown by hematoxylin–eosin and Oil Red O staining (Fig. 3B).

Transcriptomic analysis of liver tissue identified 230 genes differentially expressed between WT and Q164X heterozygotes (Fig. 3C–E). Despite the significant lipid accumulation and associated gene-expression changes in the Q164X mutant mice, we observed neither spontaneous tumor formation nor differences in Ki-67-positive hepatocyte counts compared with WT animals within the 6-month observation period (not shown).

We then extended this study by applying DEN treatment under a HFD condition to evaluate whether the metabolic alterations observed in the Q164X heterozygotes predispose them to liver tumor development. As a result, although HNF4A function was partially impaired as evidenced by reduced expression of the HNF4A target genes *HNF1A* and *ApoB* both under normal and DEN plus HFD conditions (Fig.4B), tumor formation, in both size and number, was unexpectedly more pronounced in WT mice than in Q164X heterozygous mutants. The average number of tumors in WT mice was 20.6 ± 2.0, compared to 9.0 ± 1.1 in Q164X heterozygotes. Similarly, the average tumor size was 0.97 ± 0.04 cm in WT mice and 0.36 ± 0.04 cm in Q164X heterozygotes. (Fig.4A). These findings suggest that a single-allele Q164X mutation does not promote tumor progression but may instead exert a protective effect against hepatocarcinogenesis under these experimental conditions.

**Fig. 4.**
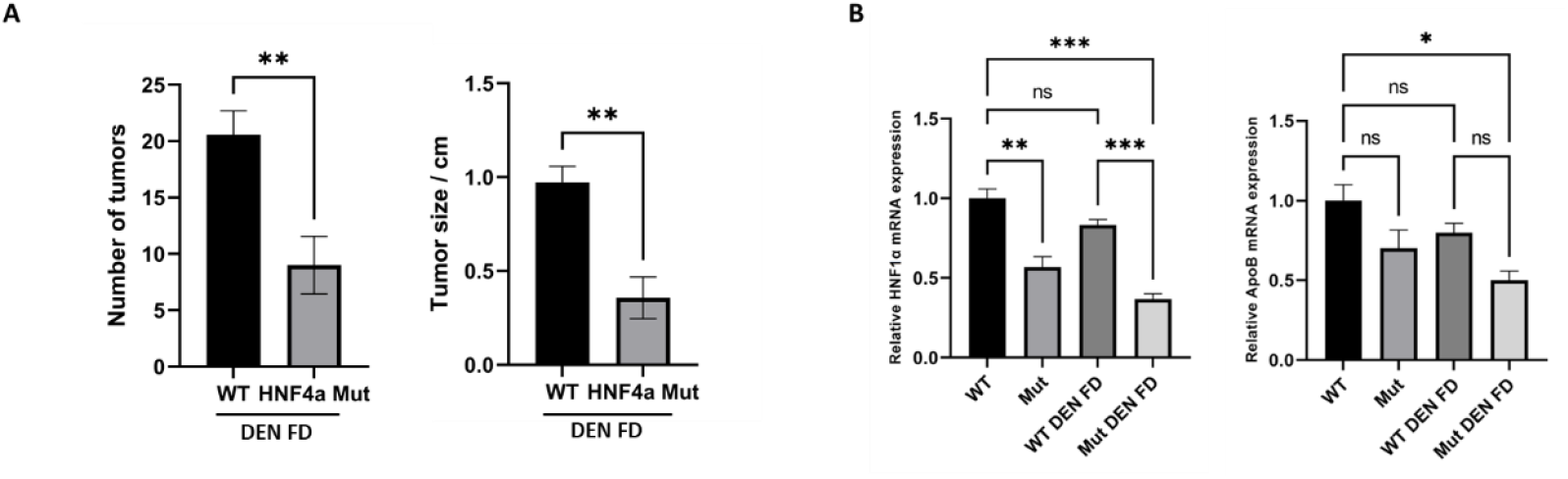
Reduced liver tumorigenesis in HNF4A Q164X heterozygous mice under DEN and high-fat diet treatment. **(A)** Quantification of liver tumor number and size in WT and HNF4A Q164X heterozygous (Q164X/WT) mice **(B)** qPCR analysis showing reduced expression of HNF4A target genes (*Hnf1a* and *ApoB*) in Q164X/WT livers compared with WT.

## Discussion

The present study provides both in vitro and in vivo evidence that the Q164X nonsense mutation in HNF4A leads to a complete loss of function and embryonic lethality when present in both alleles. At the cellular level, the Q164X mutant protein retained nuclear localization but exhibited markedly reduced transcriptional activity and DNA-binding capacity, consistent with a disruption in coactivator interactions. These results confirm that the integrity of the C-terminal region of HNF4A is essential for its transcriptional function.

In vivo, CRISPR-generated homozygous mutants were not viable, consistent with previous knockout models showing that HNF4A is indispensable for early embryogenesis and hepatocyte differentiation^19^. Heterozygous mutants survived and exhibited partial reduction of HNF4A target gene expression, including *Hnf1a* and *ApoB*, along with mild hepatic lipid accumulation, indicating compromised metabolic homeostasis. However, no spontaneous tumor formation was observed up to six months of age.

Unexpectedly, when challenged with a DEN and HFD regimen, Q164X heterozygous mice developed significantly fewer liver tumors compared with WT littermates. This paradoxical finding suggests that partial loss of HNF4A function may activate compensatory transcriptional networks that protect against carcinogenic stress. HNF4A insufficiency has been reported to shift the balance between hepatocyte differentiation and regeneration, potentially engaging factors that regulate lipid metabolism and redox homeostasis. Further transcriptomic and proteomic analyses are required to elucidate the molecular basis of this protective phenotype.

Importantly, our results indicate that structural mutations in HNF4A can cause functional inactivation independent of its expression level. While reduced HNF4A expression has been associated with hepatocellular carcinoma and poor prognosis, mutations disrupting the cofactor-binding interface may uncouple DNA binding from transcriptional activation. Thus, not all forms of HNF4A loss are equivalent structural defects and quantitative downregulation may have distinct biological consequences in hepatocarcinogenesis.

Taken together, this study identifies Q164X as a functionally null mutation in HNF4A, whose homozygosity results in embryonic lethality, while heterozygosity unexpectedly suppresses tumor formation under carcinogenic stress. These findings provide new insight into the complex relationship between HNF4A dosage, structure, and hepatic physiology, and highlight the need to investigate compensatory transcriptional and metabolic circuits that respond to partial HNF4A inactivation, particularly in human liver disorders carrying truncating HNF4A variants.

## Acknowledgements

The authors express their gratitude to Mrs. Parniansadat Khalafi and Dr. Paweł Leszczyński for their invaluable support. This research was conducted at the Department of Experimental Embryology, Institute of Genetics and Animal Biotechnology, Polish Academy of Sciences, with the facilities and resources provided by the Institute. This study was financially supported by the National Science Centre, Poland (grants no. 2017/25/B/NZ5/02762). We also acknowledge the use of ChatGPT (OpenAI) for language refinement; all suggested revisions were carefully reviewed and approved by the authors.

## Notes

The dataset is currently under preparation for submission to a peer-reviewed journal and will be modified or corrected as necessary. The English language of this manuscript was refined using ChatGPT (OpenAI), and all suggested revisions were carefully reviewed and approved by the authors.

## Notes

### Competing Interest Statement

The authors have declared no competing interest.

